# In-situ fiducial markers for 3D correlative cryo- fluorescence and FIB-SEM imaging

**DOI:** 10.1101/2021.03.14.435311

**Authors:** Nadav Scher, Katya Rechav, Perrine Paul-Gilloteaux, Ori Avinoam

**Affiliations:** Department of Biomolecular Sciences, Weizmann Institute of Science, Rehovot, Israel; Department of Chemical Research Support, Weizmann Institute of Science, Rehovot, Israel; Structure Fédérative de Recherche François Bonamy, INSERM, CNRS, Université de Nantes, Nantes, France

**Keywords:** cryo-FIB-SEM, cryo-fluorescence microscopy, Lipid droplets, correlative light and electron microscopy, multi-modal imaging, cryo-electron microscopy

## Abstract

Imaging of cells and tissues has improved significantly over the last decade. Dual-beam instruments with a focused ion beam mounted on a scanning electron microscope (FIB-SEM), which offer high-resolution 3D imaging of large volumes and fields-of-view are becoming widely used in the life sciences. FIB-SEM has most recently been implemented on fully hydrated, cryo-immobilized, biological samples. However, correlative light and electron microscopy (CLEM) workflows combining cryo-fluorescence microscopy (cryo-FM) and FIB-SEM are not yet commonly available. Here, we demonstrate that fluorescently labeled lipid droplets can serve as *in-situ* fiducial markers for correlating cryo-FM and FIB-SEM datasets, and that this approach can be used to target the acquisition of large FIB-SEM stacks spanning tens of microns under cryogenic conditions. We also show that cryo-FIB-SEM imaging is particularly informative for questions related to organelle structure and inter-organellar contacts, nuclear organization and mineral deposits in cells.

## 2. Introduction

Gaining a molecular level understanding of biological processes depends on developing the capability to visualize the molecules involved relative to their location within the cell at high resolution. Groundbreaking advances in volume electron microscopy and specimen preparation, enable the 3D visualization of cells in unprecedented detail. These advances include adapting focused ion beam milling followed by scanning electron microscopy (FIB-SEM) to the life sciences (Heymann *et al*., 2006). FIB-SEM has since been used to gain previously inaccessible insights in both cells and tissues under physiological and pathological conditions (Heymann *et al*., 2006; Knott *et al*., 2008; Merchán-Pérez *et al*., 2009; Schneider *et al*., 2010, 2011; Weiner *et al*., 2011, 2016; Reznikov *et al*., 2013; Revach *et al*., 2015). The principle of FIB-SEM is that a biological sample is exposed to a focused ion beam (usually consisting of Ga ions) capable of removing thin layers of material by milling in a highly precise manner (5-10 nm). Between each sample milling, a scanning electron beam is used to image the newly exposed surface. By repeating this process hundreds or even thousands of times, a large sample volume can be acquired with an isotropic voxel reaching 3nm (Wei *et al*., 2012). With the recent development of automated acquisition procedures, even relatively large volumes (>1000 μm^3^) can be acquired within a few days, providing large ultrastructural datasets (Peddie and Collinson, 2014; Narayan and Subramaniam, 2015). The ability to acquire large volumes with high and isometric resolution holds the potential to look at the cellular environment in a more holistic manner.

One major drawback of conventional FIB-SEM imaging at room temperatures is that it requires dehydration and resin embedding, precluding the possibility to visualize cellular structures closer to their native hydrated state (Sviben *et al*., 2016; Vidavsky, Addadi, *et al*., 2016; Vidavsky, Akiva, *et al*., 2016; Kumar *et al*., 2020). Recent studies have shown that cellular membranes are particularly visible using in-column secondary electron detection (InLens SE) in the SEM (Schertel *et al*., 2013; Spehner *et al*., 2020). This finding was unexpected because without staining by heavy atoms to generate contrast based on back-scattered electrons from the flat surface, the sample should be electron transparent. It has since been suggested that contrast may be a product of low-energy type 1 secondary electrons, generated at the electron beam focal point at low voltage (< 3kV), which are sensitive to the local surface potential of different biological material (Schertel *et al*., 2013). Nevertheless, the extent to which three dimensional (3D) organellar structure and organization can be studied at high resolution using cryo-FIB-SEM needs to be explored further.

In parallel to the increasing popularity of FIB-SEM instruments, correlative light and electron microscopy (CLEM) workflows are becoming an important tool to study rare, dynamic or undescribed cellular events by mapping information from fluorescence microscopy (FM) onto electron microscopy (EM) data of the exact same sample (Kukulski *et al*., 2011; Bykov *et al*., 2016; Karreman *et al*., 2016; Khalifa *et al*., 2016; Weiner *et al*., 2016; Weiner and Enninga, 2019; Scher and Avinoam, 2020). For correlative microscopy, visible features in both imaging modalities must be identified. This can be achieved either by using intrinsic features of the sample, such as the shape of cells, organelles and other landmarks, or by adding fiducial markers that are both fluorescent and electron opaque. The latter is typically challenging under cryogenic conditions (Masich *et al*., 2006), especially if 3D correlations of thick samples is needed, because the fiducials would have to be incorporated into the specimen volume. To overcome these challenges and develop a cryo-3D-CLEM approach, we examined whether organelles such as lipid droplets (LDs), which are relatively abundant in cells and well resolved by cryo-FIB-SEM imaging, can be used as internal fiducial markers to target the acquisition of large volumes in plunge-frozen cells grown on EM grids.

## 3. Results and Discussion

### 3.1. Organelles are well resolved in cryo-FIB-SEM

Plunge freezing of cells grown on EM grids is a well-established method to vitrify samples for cryo-EM (Medalia *et al*., 2002, 2007; Mahamid *et al*., 2016). As a first step, we grew mammalian cells on holey carbon-coated EM grids, cryo-immobilized them by plunge freezing and imaged them using mixing of InLens SE and type 2 secondary electron (SE2) detection in cryo-FIB-SEM. The SE2 detector was used to reduce the appearance of charging artifacts, caused by charge accumulation on the cross section of the sample. In all experiments (n=5), cells appeared flat on the carbon film with the region close to the nucleus accounting for most of the volume (Fig. 1A). The maximum volume acquired with an isometric voxel of 10nm was ∼5584 μm^3^, which is approximately 45% of the volume of the cell. Much like in transmission electron microscopy (TEM) membranes and lipids generated a darker contrast compared to their surroundings, as previously described. Hence, membranous organelles such as the endoplasmic reticulum, Golgi apparatus, multi-vesicular bodies, and mitochondria could be readily observed and segmented (Fig. 1B-G) (Grabenbauer *et al*., 2013; Spehner *et al*., 2020).

**Figure 1.**
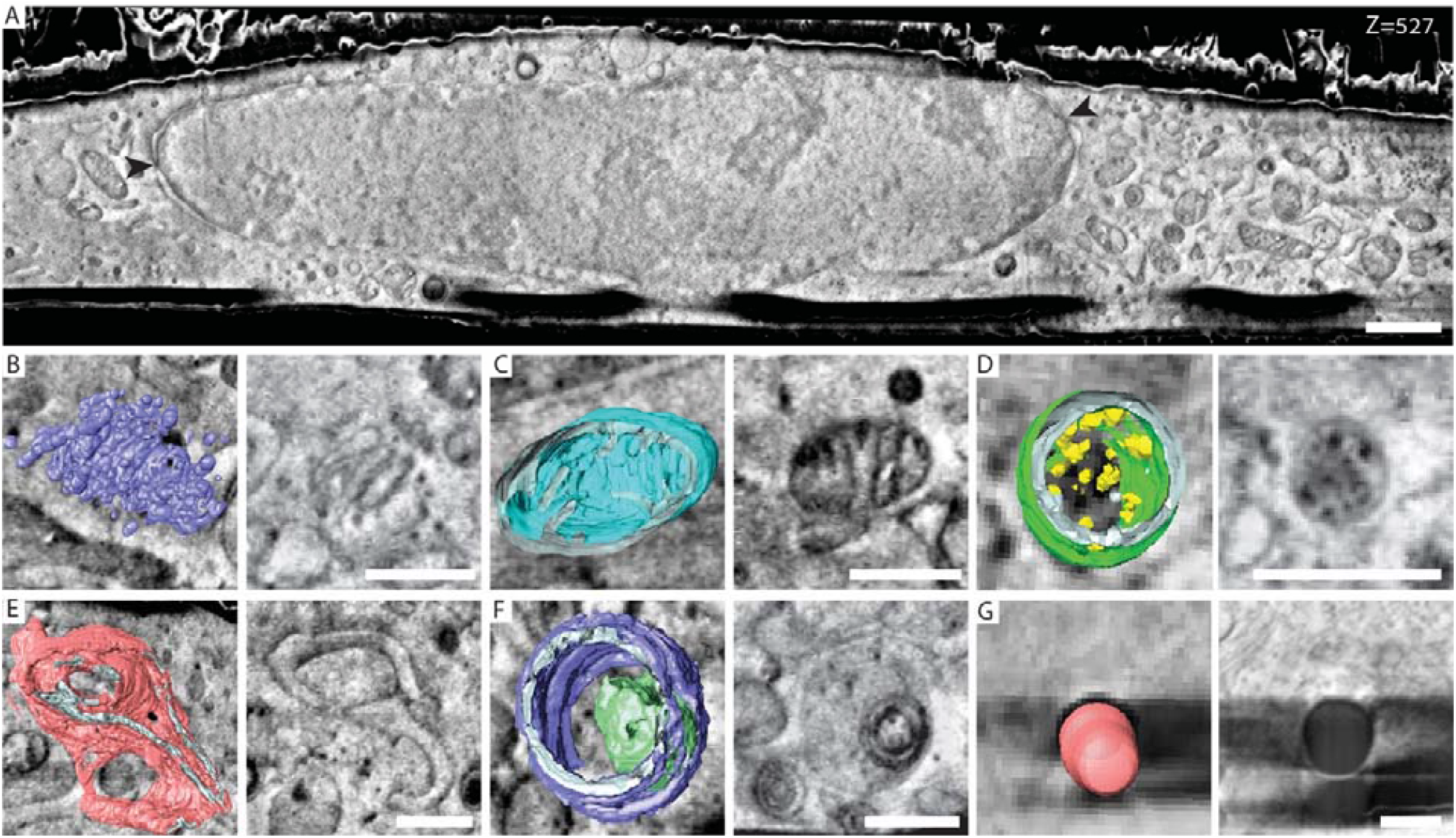
cr yo-FIB-SEM provides high resolution information of sub-cellular compar tments at their native state: **A**. A representative slice from a FIB-SEM acquisition of a plunge frozen mammalian cell. Contrast is most likely generated from differences in surface potential of the different materials. Nuclear pores (black arrowheads) Voxel size: 10×10×10 nm. Scale bar: 1 μm. **B.-G**. Overlays of representative micrographs and organelles segmentation (left) and a slice through the same organelle (right): Golgi apparatus (**B**), mitochondria (**C**), multi-vesicular body (**D**), endoplasmic reticulum (**E**), Lysosome (**F**), and lipid droplet **(G**). Scale bar: 0.5 μm.

### 3.2. Lipid droplets can serve as fiducial markers for CLEM

Re-localizing cells of interest on EM grids is relatively straight forward (Fig. 2A-C). However, targeting specific regions of interest within the huge volume of the cell is still challenging. Hence, we tested whether fluorescently labeled LDs can be used for correlative cryo-FM and FIB-SEM.

**Figure 2.**
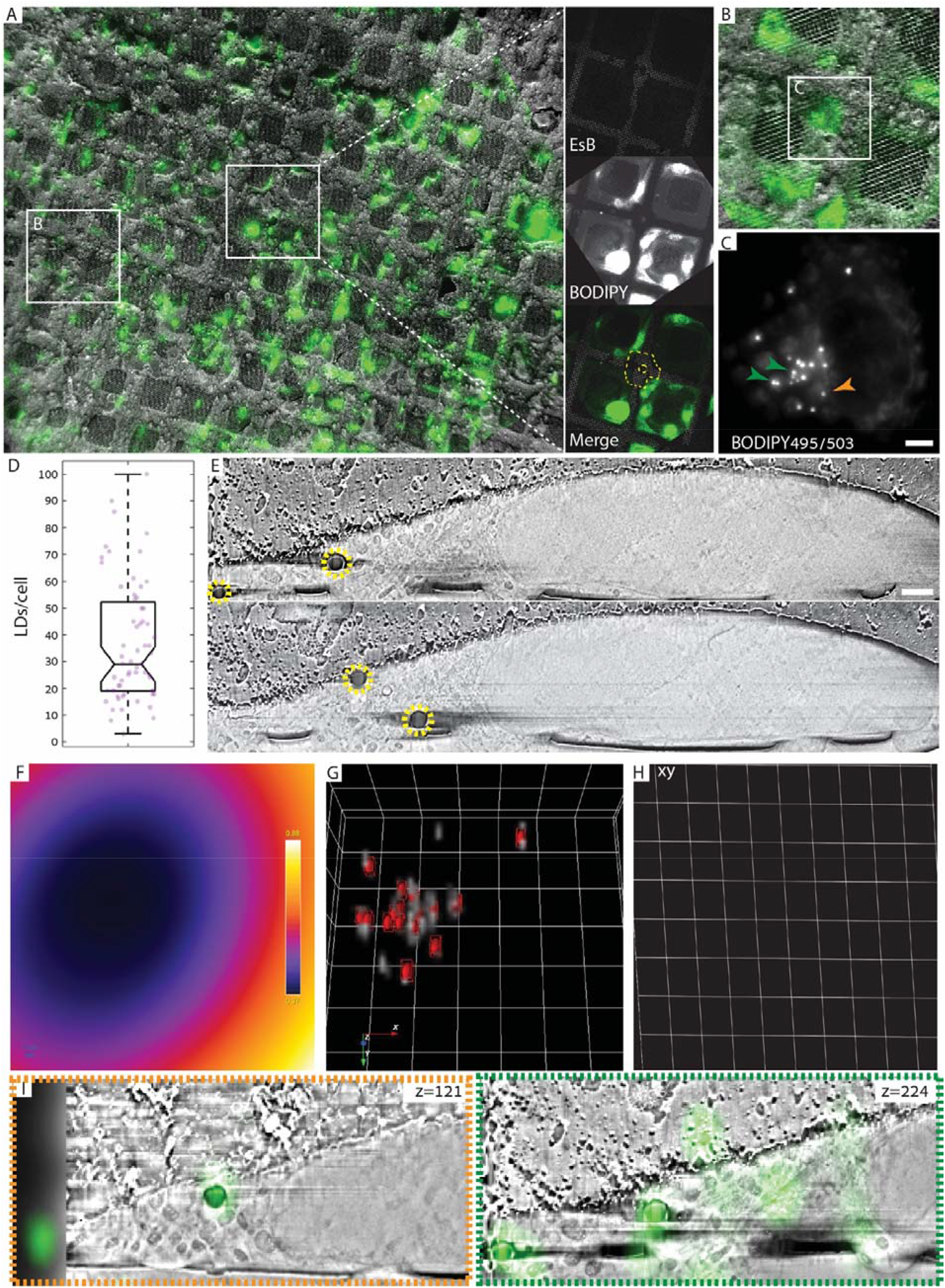
3D corr elation using lipid droplets as fiducial markers: **A**. Overlay of a low magnification cryo-FM and cryo-SEM (InLens SE / SE2 mixed detection) showing the same area on the EM grid. Cells of interest are identified in the cryo-SEM using an asymmetric mark at the center of the grid (white square and magnified on right) and other landmarks on the grid. Cells were labeled with BODIPY493/503 (green), which stains LDs. LDs are subsequently used as internal fiducial markers for CLEM. **B**. Overlay of a higher magnification cryo-FM and cryo-SEM showing a cell of interest. **C**. Maximum intensity projection of a focus series showing BODIPY distribution in a cell of interest. Arrows (green and yellow) correspond to LDs shown in the micrographs in I. Scale bar: 5 μm. **D**. Quantification of the number of LDs/cell among potential cells for CLEM. The average number is 37±22 LDs/cell (N=63, Median = 29 LDs/cell). **E**. Two representative cryo-FIB-SEM micrographs of the correlated cell, LDs (yellow circles) are clearly visible. Voxel size: 10×10×10 nm. Scale bar: 1 μm. **F**. Average z-projection of the error prediction map per image after affine image transformation using 17 LDs. The error prediction ranges from 370nm (purple) to 880nm (white). **G**. The transformed FM micrograph (gray) rendered as a volume including spheres (red) indicating the space in which the actual fiducial is with 95% confidence. **H**. Transformed grid slice of xy-plane based on the affine transformation showing the slight stretch of the image in the y and z directions (scaling of 27%, 39% respectively, z deformation is not shown). This is probably caused by artifact induced from tilt correction and line averaging (on the y axis), and minute differences in the slice thickness, and stack alignment (on the z axis). **I**. Two representative micrographs showing the overlay of the transformed FM on top of the FIB-SEM dataset. The affine transformation allowed for correlation of both LDs in the high LDs density area (green) as well as the low LDs areas (orange).

LDs, also called lipid bodies, are organelles that store neutral lipids including triglycerides and cholesterol esters (Tauchi-Sato *et al*., 2002; Olzmann and Carvalho, 2019). To use LDs as internal fiducials for correlation, we stained mammalian cells grown on grids with the fluorescent neutral lipid dye 4,4-difluoro-1,3,5,7,8-pentamethyl-4-bora-3a,4a-diaza-s-indacene (BODIPY 493/503) before plunge freezing, and visualized the cells by cryo-FM (Schorb *et al*., 2017). We observed that 100% of the cells displayed some BODIPY staining (Fig. 2), and selected cells for acquisition based on their LDs distribution. Cells suitable for acquisition showed an average of 37±22 LDs/cell (N=63, Median = 29 LDs/cell) (Fig. 2D), which assured having at least 15 LDs for correlation within a sub-volume of the cell. Whole grid landmarks such as symbols or missing grid squares were used to relocate cells in the FIB-SEM (Fig. 2A). One or two cells were acquired in one acquisition session which lasted 36-60 hours (Fig. 2E). Correlation of both datasets was performed under the ICY software (De Chaumont *et al*., 2012). We initialized the correlation by semi-automatic registration: we marked manually the centers of each LD on the FIB-SEM stack, automatically detected the centroids of LD by wavelet spot detection, and used an automatic registration on the FM dataset using the AutoFinder in eC-CLEM image registration plugin (Paul-Gilloteaux *et al*., 2017), which matches automatically the two-point sets (center of LD in FIB SEM and centroids of LD in FM) and computes and applies the rigid transformation to the image. We refined the correlation using eC-CLEM rigid transformation with 17 LDs out of 18, yielding an error estimation of a few microns (Fig. S1A). Since the rigid transformation did not give an accurate enough correlation in the particular dataset (Fig S1, S2A), we used an affine transformation on the rough registration from the Auto Finder using the same 17 LDs, yielding an error estimation between 370nm – 880nm (Fig. 2 F-I, S2C). Analyzing the resulting image transformation (Fig. 2 F-I), we observed image stretching in yz direction of FIB-SEM coordinate system (Fig. 2H). This distortion can be explained by uncompensated drift during stack acquisition. We used fast scanning with line averaging to provide an acceptable compromise between signal to noise ratio and charging artifacts. However, some charging of the surface occurs, which likely causes the slight image distortion in the xy plane. To test the robustness of the correlation with less fiducials, we repeated the refinement process using only 9 LDs, a few of which were from a cluster of LDs in the center of the cell. This approach reduced the error to 270nm-600nm, likely because minute differences in picking LDs within a tight cluster disproportionately increase the error (Fig. S2 C-D). These experiments show that LDs are suitable and attractive as internal fiducial markers for semi-automated 3D correlation.

### 3.3. Cryo-FIB-SEM as a tool to investigate cellular ultrastructure in 3D

From the volumes that we obtained in cryo-FIB-SEM, we segmented a large fraction of the mitochondrial network (Fig. 3A). We observed that interactions between adjacent mitochondria and the mitochondrial cristae could be readily resolved as well as small granules inside the mitochondria matrix (Fig. 1C, 3B). While we were unable to determine their elemental composition by energy-dispersive x-ray spectroscopy analysis (EDS) due to their small size, their size distribution suggests they are (Wolf *et al*., 2017). The Nuclear envelope (NE) and its nuclear pores were also clearly visible. The nucleus contrast was not uniform, and structures such as the heterochromatin in the periphery of the nucleus, the nucleoli, and nuclear speckles were also clearly visible (Fig. 4A). Occasionally, we observed NE invaginations into the nucleus (Fig. 4B). We segmented the NE invaginations and observed that they often followed a sinuous path (Fig. 4B). In one occasion the NE invagination formed a tube approximately 3.4 μm in length and ∼200nm in diameter that crosses the nucleus (Fig. 4C-G). We also observed small particles 42±7nm in diameter (N=5) within the NE invagination that might correspond to ribosomes (Jorgens *et al*., 2017) (Fig 4B, E-F). Together, these observations demonstrate that cryo-FIB-SEM is a particularly informative approach to study the 3D organization of membrane and membrane-less organelles in intact hydrated cells, and highlights the need to combine cryo-FIB-SEM with fluorescence information.

**Figure 3.**
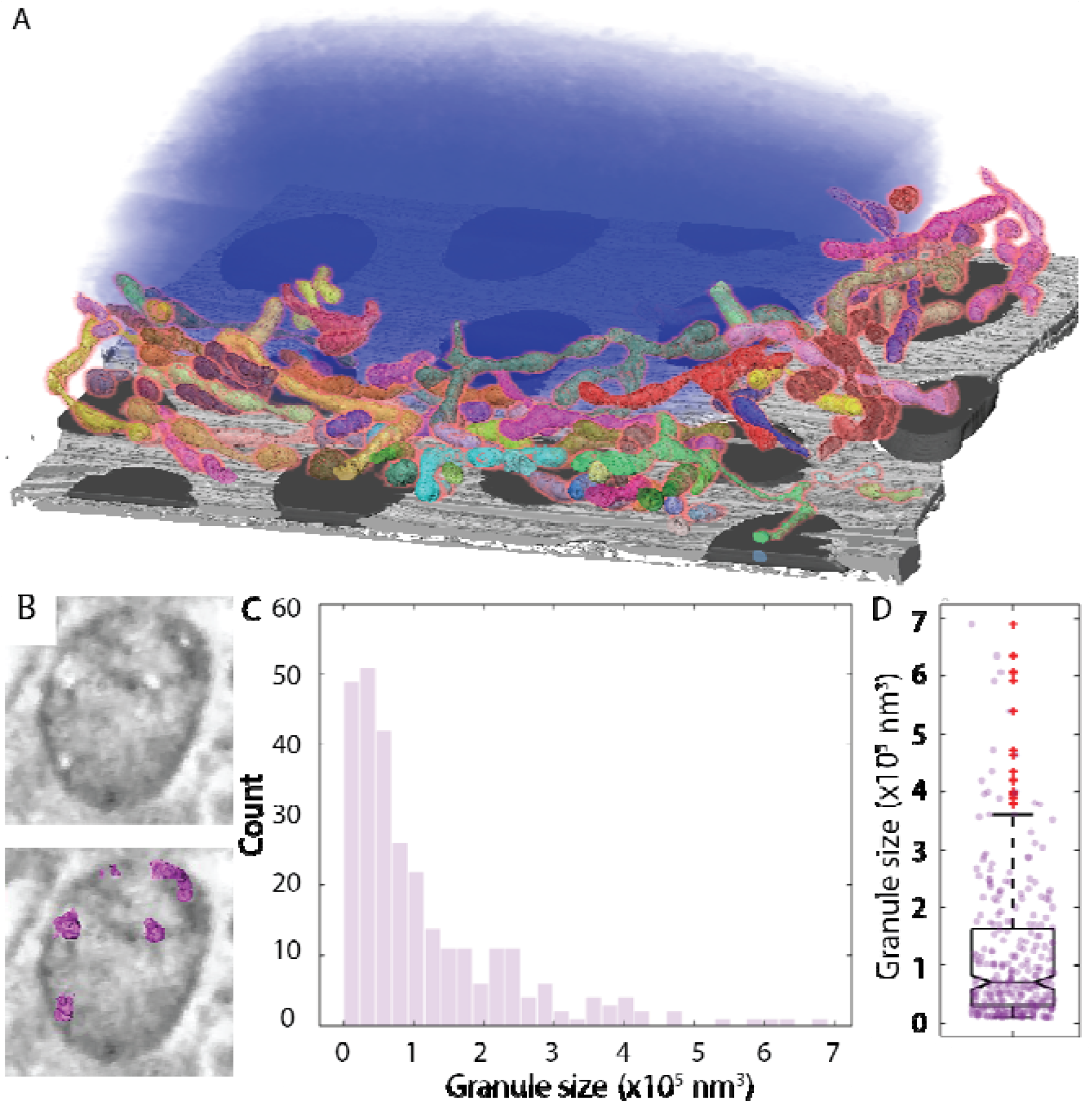
Inter- and intra-mitochondrial organization as visualized by cryo-FIB-SEM: **A**. Segmentation of part of the mitochondrial network (assorted colors). Nucleus (blue), Holey carbon support film (Gray). **B**. A cross section of one mitochondrion showing the white particles (upper image) and the same cross section with the segmented particles (magenta; bottom image). **C**. Histogram showing the size distribution of the particles inside the mitochondria matrix show similarity to size distribution of amorphous calcium phosphate granules (N=287). **D**. Box plot of the same dataset, the granules average volume is 1.16×10^5^±1.22×10^5^ nm^3^.

**Figure 4.**
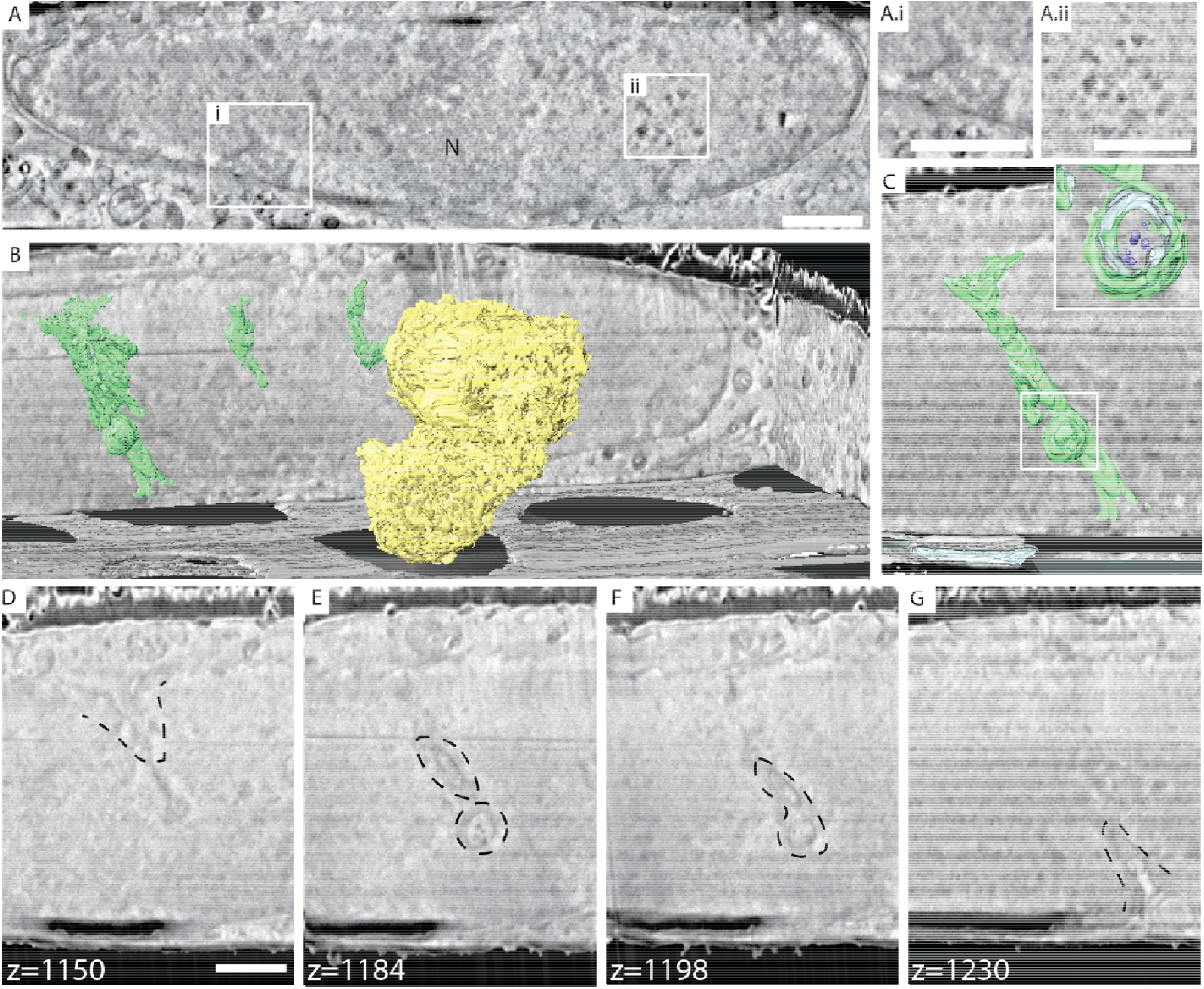
Nuclear organization as visualized by cryo-FIB-SEM: **A**. A representative slice from a cryo-FIB-SEM acquisition highlighting the nucleus. Differences in nuclear content is visible based on contrast change. N-nucleolus, **A.i**-heterochromatin, **A.ii**-nuclear speckles. Scale bar: 1 μm. **B**. Segmentation of nucleoli (yellow) and nuclear envelope invaginations (green). One of the observed invaginations passed through the whole nucleus and was less sinuous then others. Holey carbon support film (Gray). **C**. The nuclear invagination crossing the entire nucleus spanning approximately 3.4 μm long, and approximately 200nm in diameter. Inset shows the particles inside the invagination (blue). **D-G**. Representative slices through the nuclear envelope invagination showing the particles inside the spherical appendage (**E-F**). The small particles measured are 42±7nm in diameter (N=5), and likely correspond to ribosomes. Scale bar: 1 μm.

## 4. Conclusions

The Cryo-FM and FIB-SEM correlative imaging workflow presented here can be used to study any cell that has LDs dispersed around the region of interest. It is particularly useful for studying cellular structures that are not well preserved by conventional FIB-SEM. The resolution and correlation precision can be further improved by using cryo-confocal or super-resolution FM (Arnold *et al*., 2016; Wolff *et al*., 2016; Hoffman *et al*., 2020). Improving the correlation precision may be useful if the CLEM workflow is used for cutting thin lamella for TEM rather than for FIB-SEM imaging. Much can also still be done to enhance the success rate of data acquisition, alignment, image processing and analysis.

The workflow will be particularly useful for pinpointing specific regions of the mitochondrial network, unambiguously identifying organelles or sub-structures within the nucleolus, in order to resolve their underlying ultrastructure in 3D. It would also be beneficial for identifying specific cells or sub-cellular regions in more complex samples such as tissues. Cryo-FIB-SEM imaging appears highly suited for questions related to organelle remodeling and nuclear organization because it provides sufficient resolution and a more complete view of larger objects than what can often be gauged from thin sections.

In conclusion, we demonstrated the use of organelles as fiducial markers for cryo-CLEM as an alternative for external fiducial markers to the specimen. Specifically, we show the usefulness of LDs as fiducials for correlation. While we used BODIPY staining for this study, other neutral lipid dyes in the red and far-red wavelengths can be used. In combination with fluorescent reporters highlighting specific transient processes or states, this workflow can be used to address different questions in cell biology where resolving the ultrastructural organization of the cell at its native state is important. Since cryo-confocal and super-resolution FM have already been realized and developed, they can be applied when higher resolution or thicker high-pressure-frozen samples are needed. Cryo-FIB-SEM tomography is clearly a promising imaging approach that is not commonly used.

## 5. Acknowledgements

We thank Dr. Luca Bertinetti for providing scripts for image processing, Dr. Martin Schorb from the EMBL electron microcopy core facility for assistance in optimizing the cryo FM image acquisition, the EM unit of the Weizmann Institute of Science, Dr. Neta Varsano and Dr. Mattia Morandi for critical reading of the manuscript, and members of the Avinoam team for fruitful discussions. P.P-G is part of MicroPICell facility (BioGenouest), member of the national infrastructure France-BioImaging (ANR-10-INBS-04), and acknowledges funding from ANR-18-CE45-0015. This research was supported by the Minerva Foundation with funding from the Federal German Ministry for Education and Research, the David Barton Center for Research on the Chemistry of Life. This project has also received funding from the European Research Council (ERC) under the European Union’s Horizon 2020 research and innovation programme (grant agreement No 851080). O.A is the Miriam Berman Presidential Development Chair.

## 6. Authors contribution

N.S and O.A conceived and designed the experiments and analyzed the data. N.S with help from K.R carried out the experiments. N.S with help of P.P-G correlated the data. N.S and O.A wrote the manuscript.

