## Supplementary for "In-situ fiducial markers for 3D correlative cryo- fluorescence and FIB-SEM imaging"

### 1. Supplementary figures

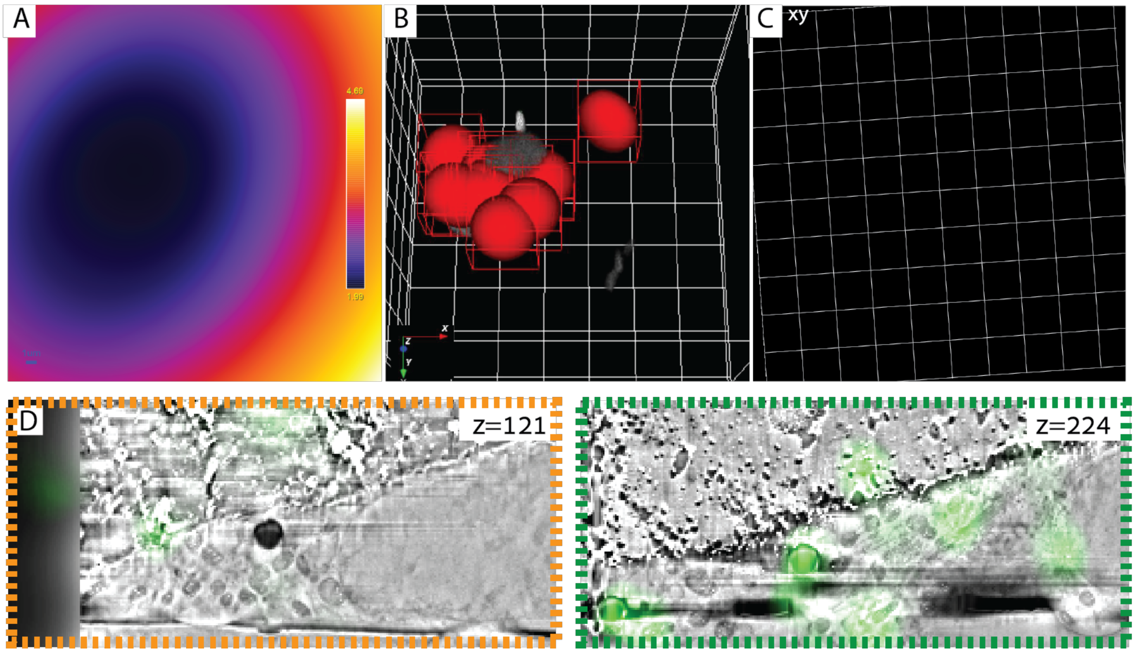


**Supplementary Figure 1. 3D rigid correlation using 17 LDs: A.** Average projection of the error prediction map across the stack. The error in rigid transformation ranged between 1.99µm (purple) and 4.69µm (white). **B.** The transformed FM micrograph (gray) with spheres representing the most probable area to find each fiducial (red) with 95% confidence interval. The spheres volume is far greater compared to the affine correlation presented in Fig. 2G. **C.** Transformed xy-plane grid image showing the rigid transformation result. The squares are even and are not stretched as in the affine transformed grid in Fig. 2H. **D.** Representative slices showing the overlay in transformed FM micrographs and FIB-SEM micrographs. While in the middle of the stack the correlation is decent (green box) in the edges of the dataset there is a large discrepancy (orange box).

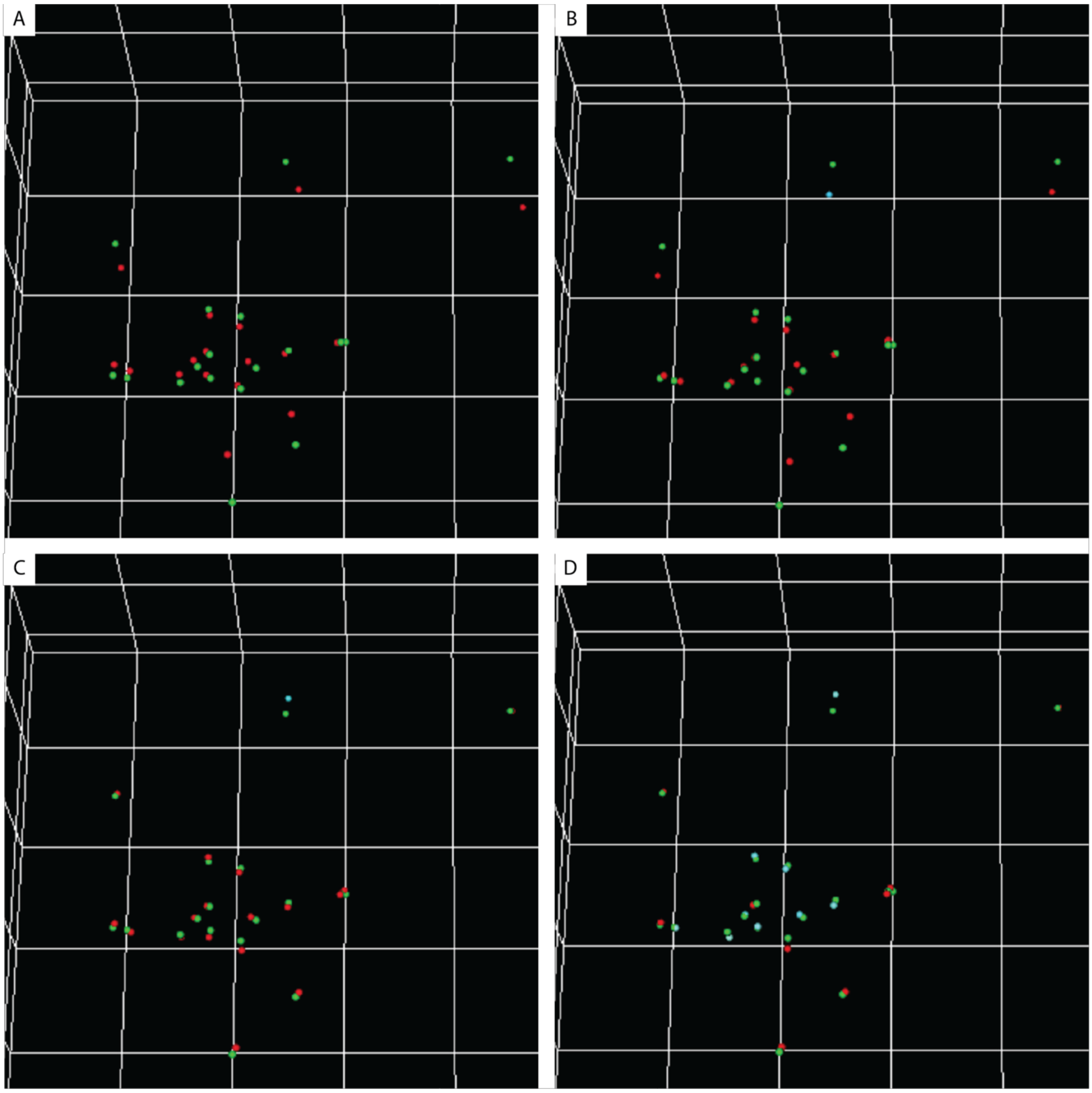


**Supplementary Figure 2. Distances between LDs after different image registration approaches:** Distance between LDs after different registrations shown in each panel, where green spheres are the LDs position in the FIB-SEM dataset, red spheres are the LDs position in the fluorescence dataset after registration used to compute the transformation, and cyan sphere are the LDs position in the fluorescence dataset after registration, but which were not used for the registration. **A.** after the automatic matching by the autofinder constraining the transformation to be rigid (no scaling allowed). **B.** after the autofinder, refined registration by manual matching using 17 lipid droplets (red) and constraining the transformation to be rigid; in which the average distance between the same fiducial in EM and FM was 1085.5±1086.1nm. **C.** after the autofinder, refined registration by manual matching using 17 lipid droplets (red) and constraining the transformation to be affine (scale shearing allowed); **D.** after the autofinder, refined registration by manual matching using 9 lipid droplets (red) and constraining the transformation to be affine (scale shearing allowed). The average distance between the same fiducial in EM and FM improved significantly to 220.3±164.0nm One grid square is 5 microns in every axis.

### 2. Materials and methods

2.1. Cell culture and staining

Murine C2C12, were cultured in growth medium (GM) composed of modified Dulbecco's Modified Eagle Medium (Gibco) supplemented with 10% FBS (Gibco), 1% sodium pyruvate (Biological industries), 1% penicillin-streptomycin (Biological industries) and 20mM HEPES (Biological industries), and maintained at 37°C, 5% $CO_2$. For Cryo-FM, cells were seeded on Quantifoil R3.5/1 Au 200 mesh EM grids (Quantifoil Micro Tools, Germany) at a cell density of 340 cells/$mm^2$. After 18-24 hours cells were stained with BODIPY 493/503 (1:1000 in DPBS; ThermoFisher Scientific, Waltham, MA, USA) at 37°C for 20 min followed by three washes in GM.

2.2. Cryo-immobilization and cryo-FM

Grids were plunge-frozen between 45-90 min after staining using a Leica EM GP (Leica Microsystems, Vienna, Austria) set to 37°C, 90% humidity and following 2.5s back-blotting. To facilitate their handling, grids were loaded into c-clip rings (Thermo Fisher Scientific) under liquid nitrogen prior to imaging. Imaging was performed using a cryo-CLEM microscope (Leica Microsystems) as previously described (Schorb *et al.*, 2017). Samples were mounted onto a costume-made cartridge (EMBL, Heidelberg, Germany) in a liquid $N_2$-cooled cryo-CLEM shuttle and transferred to the microscope. Samples were imaged under HCX PL APO 50× CLEM cryo-objective and images were acquired using an ORCA-flash4.0 (Hamamatsu Photonics, Hamamatsu city, Japan) with GFP (ex450-490/em500-550) filter. Initially, a spiral scan was acquired in bright-field (50ms exposure, 100% lamp intensity). To screen the grid and identify cells of interest, a grid map was by acquiring (250ms exposure, 100% lamp intensity) and stitching images with step size of 1-2μm over 50-100μm. We chose between seven and fifteen cells per grid based on their LD signal. For cells of interest, focus series were acquired using 25ms exposure, (brightfield) and 200ms exposure (GFP), and a step size between 150-250nm over a 10μm range.

2.3. Cryo-FIB-SEM

Grid in c-clip rings were mounted onto a costume-made cryo-sample holder inside a modified Leica EM-VCM500 (Leica Microsystems, Vienna, Austria) and transferred to a pre-cooled Zeiss Crossbeam 550 FIB-SEM (-150°C, Carl Zeiss Microscopy, Oberkochen, Germany) using Leica EM-VCT100 shuttle (Leica Microsystems, Vienna,

Austria). The Leica EM-VCM500 is closed to form a glove box and dried using constant flow of clean $N_2$ gas in order to reduce exposure to humidity. The cryo-sample holder has a pre-tilt of -25° and the cryo-stage was tilted to 35°-40° during acquisition. Pt precursor deposition was performed by opening and closing the nuzzle of the multi-port gas-injection system in 3 cycles of 30s, without heating the source, while working distance was 7-8mm. FIB milling was executed by SmartFIB software (Carl Zeiss) using 30kV and probe current of 100pA to cut 10nm-thick sections, ion beam dose factor was 8, and dwell time 160µs. SEM micrographs were acquired at 1.6kV acceleration voltage with a probe current of 40pA and InLens SE/SE2 mixed detection. Scanning speed of SEM was 1 and noise reduction was performed by line averaging (Fig. 1, 3-4: N=104, Fig 2: N=140). The voxels sizes in data showed are 10x10x10 nm. For overnight acquisition, Liquid $N_2$ was replenished using an automatic liquid $N_2$ handling system (Norhof LN2 microdosing systems, Ede, The Netherlands).

### 2.4. Image Processing and Segmentation

Images were filtered using Fourier filtering approach with either Fiji (Schindelin *et al.*, 2012) or a Fourier filtering Python 3.7 script kindly provided by Dr. Luca Bertinetti in order to reduce curtaining artifacts (Spehner *et al.*, 2020). Stack alignment was performed either by cross-correlation alignment using Align Slices module in Amira 2019.3 (Thermo Fisher Scientific) or based on Fourier shift theorem as described in Spehner *et al.* (Spehner *et al.*, 2020), Python 3.7 script was kindly provided by Dr. Luca Bertinetti. Both alignment methods allowed only for translation of the slices and not rotation or scaling. To enhance contrast, unsharp mask filter was applied to the stacks in Fiji software (Schindelin *et al.*, 2012). In order to reduce charging artefact, charge correction was performed as described in Spehner *et al.* with a Python 3.7 script kindly provided by by Dr. Luca Bertinetti (Spehner *et al.*, 2020).

Data segmentation was performed manually using Amira 2019.3 using the threshold tool on every 3$^{rd}$ or 4$^{th}$ slice. The rest of the slices were interpolated. Intra mitochondrial granules were segmented without interpolation, owing of their small size. All the segmented labels were converted into surfaces using Generate Surface module and presented using a Surface View object. To represent the volume of the nucleus in Fig. 3, volume rendering was applied to the whole stack and the excess volume was cropped out using Volume Edit module.

### 2.5. Quantifications and Correlation

Intra-mitochondrial granules quantification was performed based on the segmented particles. Volume of each particle was calculated by Amira using Material Statistics feature per particle. To avoid manual errors in the segmentation, object smaller than 10 voxels were not considered. The histogram and box plot chart were plotted using MATLAB 2019b (MathWorks).

LDs count per cell was made using the following processing steps in Fiji (Schindelin *et al.*, 2012): Cells of interest were cropped and an intensity z-projection was calculated based on standard deviation. The image was then threshold to form a binary image and using the watershed tool. The LDs were counted using the analyze particles tool and corrected manually for false positive and false negatives. Box plot was plotted using MATLAB 2019b.

For 3D correlation, the processed cryo-FIB-SEM was binned in Fiji (Schindelin *et al.*, 2012). The center off mass was manually picked in the FIB-SEM dataset and the BODIPY signal was segmented using spot detector (parameters: detect bright spots, scale 2, sensitivity 10, export to ROI) in Icy (De Chaumont *et al.*, 2012). Rough, rigid transformation was applied using Auto Finder feature in eC-CLEM (Paul-Gilloteaux *et al.*, 2017) (parameters: FM with detected ROI spots as source image, FIB-SEM with manually picked ROI as target image, pre-aligned data, max error 10 microns, percentage of point to kept 70%, “already in the same orientation” option) following by either rigid or affine transformation with either 9 or 17 LDs identified manually under ec-CLEM v2.1.0. Error estimation heatmaps and 95% interval confidence spheres where calculated by eC-CLEM v2.1.0.
